# Revisiting a classic hybrid zone: rapid movement of the northern flicker hybrid zone in contemporary times

**DOI:** 10.1101/2021.08.16.456504

**Authors:** Stepfanie M. Aguillon, Vanya G. Rohwer

**Affiliations:** Department of Ecology and Evolutionary Biology, Cornell University, Ithaca, NY 14853; Fuller Evolutionary Biology Program, Cornell Lab of Ornithology, Ithaca, NY 14850; Cornell University Museum of Vertebrates, Ithaca, NY 14850

## Abstract

Natural hybrid zones have provided important insights into the evolutionary process, and their geographic stability/instability over time can help to disentangle the underlying biological processes that maintain them. Here, we leverage replicated sampling of an identical transect across the hybrid zone between yellow-shafted and red-shafted flickers in the Great Plains to assess its stability over ∼60 years (1955-1957 to 2016-2018). We identify a ∼73 km westward shift in the hybrid zone center towards the range of the red-shafted flicker, but find no associated changes in width over our sampling period. In fact, the hybrid zone remains remarkably narrow, suggesting some kind of selective pressure maintains the zone. By comparing to previous work in the same geographic region, it appears likely that the movement in the hybrid zone has occurred rapidly in the years since the early 1980s. This recent, rapid movement may be related to changes in climate or land management practices that have allowed asymmetric westward movement of yellow-shafted flickers into the Great Plains.

## Introduction

Naturally hybridizing taxa provide unique insights into the process of speciation (Barton and Hewitt 1985; Harrison 1993; Harrison and Larson 2014). Hybrid zones, geographical regions where differentiated taxa interbreed and produce hybrids, have long-been described as “windows on evolutionary process” as they provide opportunities to assess the outcome of hybridization over many generations (Harrison 1990). Additionally, the geographic locations of hybrid zones can provide important insights and movement of hybrid zones has been of particular interest in recent years (e.g., as “windows on climate change” Taylor et al. 2015). Molecular methods have recently made it possible to identify signatures of historical hybrid zone movement in the genome (Wielstra et al. 2017; van Riemsdijk et al. 2019; Wielstra 2019), but this inferred evidence of movement does not always match results from direct resampling over broad temporal and spatial scales (Wang et al. 2019). Although difficult to accomplish, repeated sampling of hybrid zones over time remains the best way to definitively identify movement.

Repeated sampling is additionally important because hybrid zone movement can occur under models where hybrids have lower or higher fitness than parental taxa. When hybrids have lower fitness—the tension zone model—a balance between selection against hybrids and parental dispersal into the hybrid zone influences the location of the hybrid zone (Barton and Hewitt 1985, 1989). Differences in population density of the hybridizing taxa (Barton and Hewitt 1985), asymmetric hybridization (Konishi and Takata 2004), or differential adaptation of parentals (Key 1968) could lead to hybrid zone movement. On the other hand, when hybrids have higher fitness—as in an environmental selection gradient model—the hybrid zone is often bounded within an intermediate environment (Moore 1977). Competitive advantage of one taxa over the other (Buggs 2007), asymmetric hybridization (Buggs 2007), or changes in the environment (Taylor et al. 2015) could lead to hybrid zone movement. Repeated sampling can help to disentangle the mechanisms that are maintaining the hybrid zone itself.

Here, we directly assess movement in the hybrid zone between yellow-shafted (*Colaptes auratus auratus*) and red-shafted (*C. a. cafer*^1^) flickers. The flicker hybrid zone is a long-studied system in ecology and evolution (e.g., Short 1965; Moore and Buchanan 1985; Wiebe 2000) that has intrigued naturalists since at least the mid-1800s (Audubon et al. 1897). Flickers are common woodpeckers widely distributed across wooded areas of North America—red-shafted flickers in the west and yellow-shafted flickers in the east (Wiebe and Moore 2020). These two forms come into secondary contact in an extensive hybrid zone in the Great Plains that roughly follows the Rocky Mountains from northern Texas to southern Alaska (Fig. 1). Hybridization between the flickers is clearly visible due to differences across six distinct plumage traits (Fig. 1, Table S1), yet there is only mixed evidence of assortative mating based on these traits—with no evidence in the US portion of the hybrid zone (Bock 1971; Moore 1987) and weak but significant evidence in the Canadian portion of the hybrid zone (Wiebe 2000; Flockhart and Wiebe 2007; Wiebe and Vitousek 2015). To date, no negative fitness consequences of hybridization have been identified in any part of the hybrid zone (Moore and Koenig 1986; Wiebe and Bortolotti 2002; Flockhart and Wiebe 2009), and it has been hypothesized that hybrids actually have higher fitness than parentals (Moore and Price 1993). Previous work found stability in both the center and width of the hybrid zone in the US up to the early 1980s (Moore and Buchanan 1985).

**FIGURE 1.**
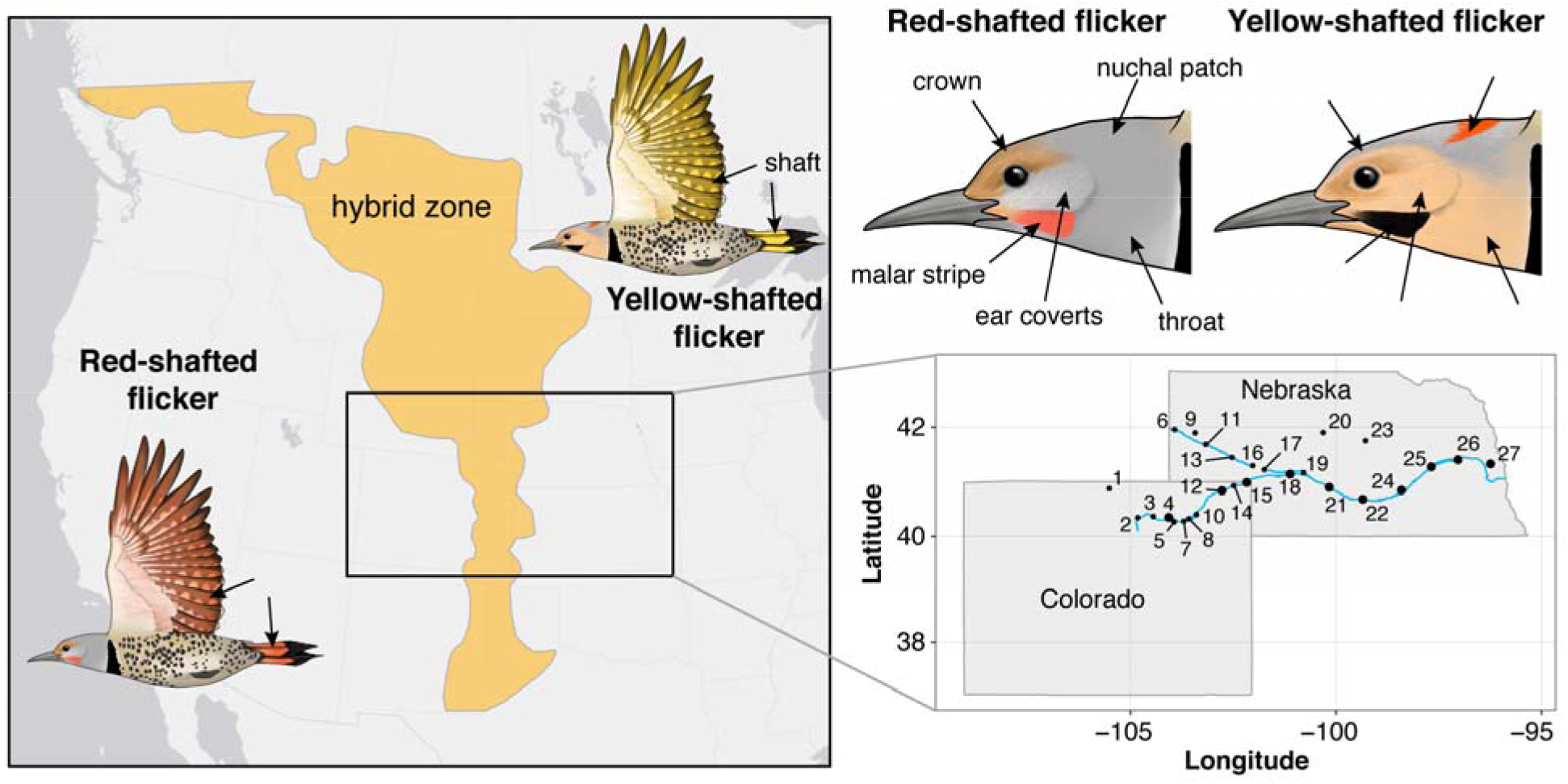
The geographic extent of the southern portion of the hybrid zone between red-shafted and yellow-shafted flickers as estimated in Moore and Price (1993). The inset map of Colorado and Nebraska depicts the repeated sampling conducted along the Platte River with numbered sampling localities (Table S3). Larger points on the inset map indicate localities that were sampled during both the historic (1955-1957) and contemporary (2016-2018) sampling periods. The six plumage differences are shown with arrows on the flicker illustrations (created by M. Bishop) and details are described in Table S1.

Here, we assess movement of the flicker hybrid zone in the US Great Plains by comparing the spatial transition of plumage characters between two sampling periods of an identical transect separated by ∼60 years (Fig. 1 inset). Evaluating patterns in plumage traits is a powerful approach to understand hybrid zone dynamics at both the phenotypic and genotypic level in flickers, as they have extremely low levels of genomic divergence (Aguillon et al. 2018) with the few existing differentiated regions being associated with plumage color differences (Aguillon et al. 2021). We first validate our plumage scoring approach using independent multispectral photography. Using geographic cline analyses, we then estimate the center and width of the plumage clines in the historic (1955-1957) and contemporary (2016-2018) sampling periods and compare them. Finally, we evaluate changes in the cline center and width against changes expected under a model of neutral diffusion (Wang et al. 2019).

## Material and Methods

### SAMPLING

The most extensive study of hybridization and phenotypic variation in flickers was undertaken by Short (1965), for which he collected specimens intensively along the Platte River in Nebraska and Colorado from 1955-1957 during the breeding season (Fig. 1 inset). This hybrid zone transect for flickers (and those for several other hybridizing species pairs) remains one of the most extraordinary components of the ornithological collection in the Cornell University Museum of Vertebrates (CUMV). During the spring and summer of 2016-2018, the CUMV replicated Short’s sampling along the Platte River—revisiting many of his original localities—to amass a modern-day transect of the hybrid zone. This was additionally supplemented by banding individuals in 2016.

Henceforth, we will use “historic” and “contemporary” to refer to flickers sampled from 1955-1957 and 2016-2018, respectively. We focus here on adults to avoid confounding patterns due to immature plumage in juveniles. We include 252 historic flickers (all vouchered in the CUMV) and 107 contemporary flickers (91 specimens vouchered in the CUMV and 16 individuals that were banded, photographed, and released) in this study (Table S2). We group the sexes together across all analyses except for those on the malar stripe (the only sexually dimorphic character in flickers), where we include only males (138 historic and 72 contemporary).

### PLUMAGE SCORING

The flickers differ across six primary plumage characteristics (Fig. 1): the eponymous “shaft” (wing and tail) color, crown color, ear covert color, throat color, malar stripe color in males, and the presence/absence of the nuchal patch (Short 1965). In brief, these birds differ vividly in the shaft color (bright yellow in the yellow-shafted flicker versus salmon red in the red-shafted flicker) and in the overall coloring of the face and head. Hybrids can exhibit various combinations of the six parental traits, as well as colors intermediate to the parental extremes.

We scored plumage characters of historic and contemporary flickers on a categorical scale from 0 (pure yellow-shafted) to 4 (pure red-shafted) for each of the six plumage traits following a protocol slightly modified from Short (1965; see Fig. 1 and Table S1 for details); a method that has been used extensively within the flicker system (e.g., Moore and Buchanan 1985; Moore 1987; Wiebe 2000; Flockhart and Wiebe 2007; Aguillon et al. 2021). The main modification from Short (1965) is differences in our scoring of the shaft color based on an increased understanding of carotenoid pigmentation, particularly around orange shaft feathers (e.g., Hudon et al. 2017). We additionally calculated an overall plumage hybrid index by summing across the trait scores within each individual and dividing by 24 in males and 20 in females (the maximum possible) to obtain a score that ranges from 0 to 1. This standardization makes comparisons between males and females possible, as females lack the malar stripe present in males. All scoring was conducted by the first author to ensure consistency.

### COMPARISON OF PLUMAGE SCORING WITH MULTISPECTRAL PHOTOGRAPHY

To assess the accuracy of our plumage scoring method, we collected multispectral images of all flickers in the contemporary sampling period. For complete details on image collection and processing, see Supplemental Text S1 and Ligon et al. 2018. In brief, we photographed each specimen from three viewing angles (ventral, dorsal, and lateral) and under two conditions (all visible light between 400-700 nm and UV light between 300-400 nm). We then created standardized multispectral image files using the micaToolbox (Troscianko and Stevens 2015) in ImageJ (Schneider et al. 2012), and outputted values for each color channel (red, green, blue, UV) and luminance for each of the six plumage traits, as well as the overall area for the nuchal patch. We performed multiple regressions to directly compare the plumage score with the image parameter values obtained from the multispectral photography. For the crown, ear coverts, malar stripe, shaft, and throat we compared the plumage score to the values for the four color channels and luminance. For the nuchal patch we compared the plumage score to the area.

### GEOGRAPHIC CLINE ANALYSES AND HYBRID ZONE MOVEMENT

To evaluate the distribution of phenotypic traits across the hybrid zone, we fit a series of equilibrium sigmoidal cline models (Szymura and Barton 1986; Gay et al. 2008) using the ‘nls’ function in R v.3.6.2 (R Core Team 2018), where we modelled the relationship between locality (*x*) and hybrid index or trait score (*y*) to estimate cline center and width. *S* is included as a scaling factor for trait scores that do not vary from 0 to 1. Confidence intervals for center and width were calculated using the ‘confint’ function in R. We repeated this process for both historic and contemporary sampling periods and compared the results for the overall hybrid index (*S*=1) and individual phenotypic traits (*S*=4).

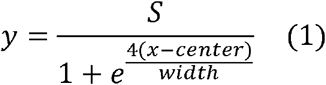

We grouped samples based on sampling location (Fig. 1, Table S3) and set the start of the cline to the western-most locality sampled in the Rocky Mountains. To estimate the distance of each locality from the start of the cline, we determined the mean latitude across all localities and then used the ‘distm’ function in the geosphere R package (Hijmans 2019) to calculate the distance each locality was from the start of the cline along the mean latitude value, using the longitude of the locality and assuming an ellipsoid shape.

We additionally assessed changes in the hybrid zone between the two sampling points using the overall hybrid index in two ways following an approach taken by Wang et al. (2019). First, to assess movement of the hybrid zone center, we used AIC to compare the contemporary cline to a model using the contemporary data but with the cline center fixed on the estimated historic center. This approach accounts for uncertainty due to sampling error between the two sampling periods. If there was no difference between the estimated cline centers for the two time periods, the fixed center model would be expected to have lower AIC than the true model. Second, to assess change in the width of the hybrid zone and see if selection is maintaining the cline, we followed an approach developed by Wang et al. (2019) that leverages repeated sampling of the same transect over time to test against the neutral diffusion model (Barton and Hewitt 1985).

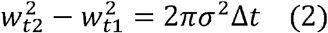

Where 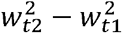 is the difference in cline width squares between the two sampling points, *σ* is a measure of dispersal distance, and Δ*t* is the number of generations between the sampling points. We calculated the bootstrap distribution of 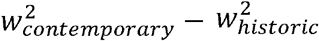 by resampling with replacement and fitting clines over 100,000 iterations.

We used both realistic and conservative values to estimate the neutral diffusion expectations from equation (2). For more realistic values, we used *σ* a of 100.7 km as estimated by Moore and Buchanan (1985) using banding data and a Δ*t* of 60 generations (i.e., 1 year/generation). For more conservative values, we used a *σ* of 30 km as natal dispersal is typically greater than 15 km (and likely much greater; Wiebe and Moore 2020) and a Δ*t* of ∼33.3 generations (i.e., 1.8 years/generation; Milá et al. 2007).

## Results

We validated our categorical scoring approach of flicker plumage coloration using multispectral photography. Multiple regressions comparing the plumage scores with the image parameter values were strongly significant for all six plumage traits (Table S4, S5; Fig. S1).

We detected a significant westward shift of ∼73 km in the hybrid zone cline center between the historic and contemporary sampling periods for the plumage hybrid index (Fig. 2A, Table S6). The individual clines for the six plumage traits are broadly overlapping within both the historic (Fig. S2A) and contemporary (Fig. S2B) periods. The cline center from the historic sampling period did not fit the contemporary data (Fig. 3A; AIC = -32.7 for the true model, AIC = -6.6 for the model with the historic center), further supporting the movement of the hybrid zone between the two sampling periods. However, we did not identify associated changes in hybrid zone width between the historic and contemporary sampling periods (Table S6). In fact, the bootstrap distribution of 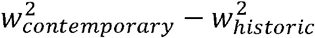 (95% CI: -37,441, 90,116) was significantly less than predicted by the neutral diffusion model under both realistic 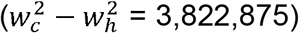 and more conservative 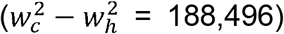 dispersal distance and generation time values (Fig. 3B), suggesting that selection has maintained the narrow width of the hybrid zone.

**FIGURE 2.**
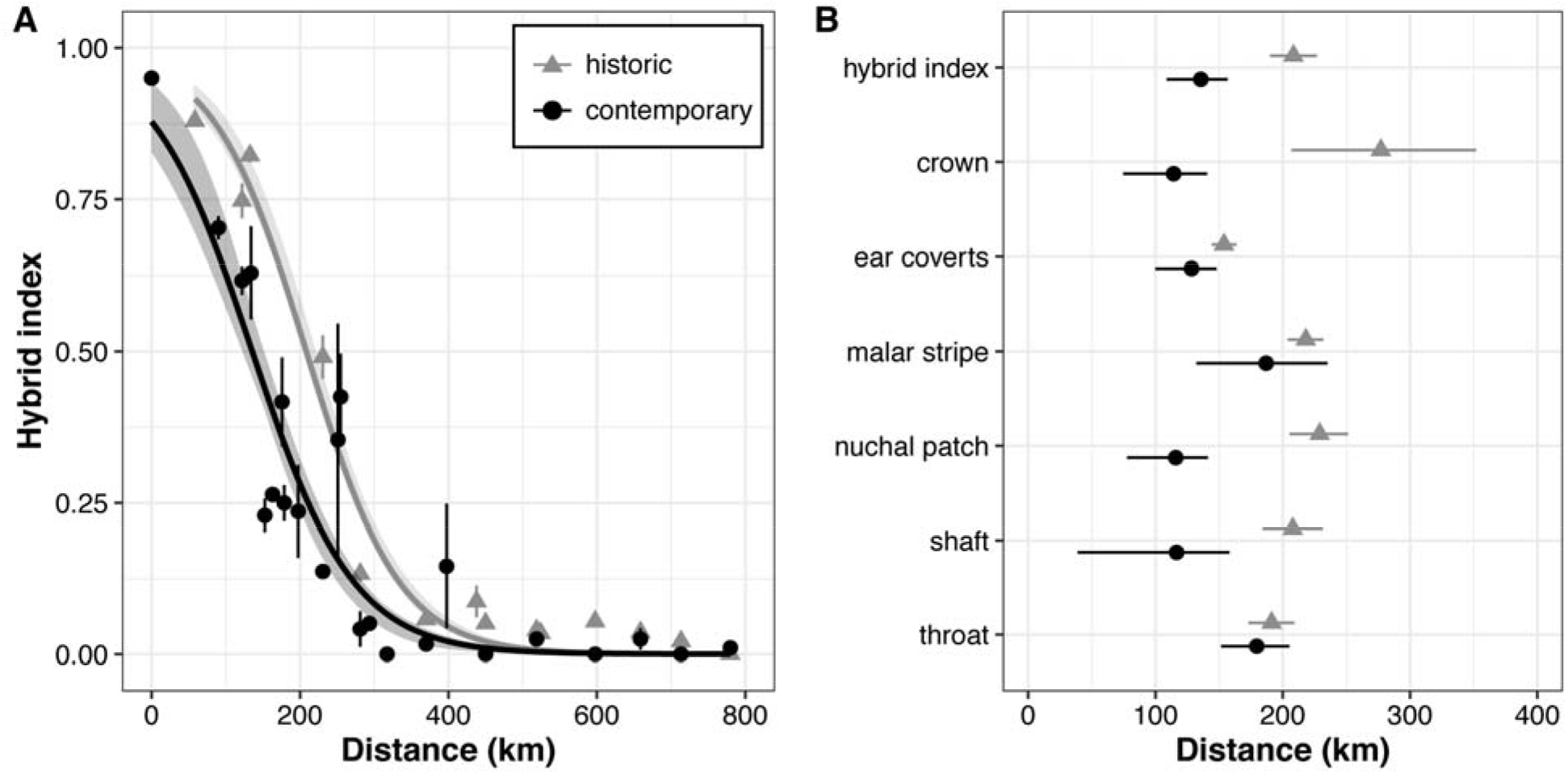
(**A**) Geographic clines of the overall hybrid index as estimated for historic (gray, triangles) and contemporary (black, circles) flickers demonstrate the ∼73 km westward movement of the hybrid zone in the ∼60 years between the two sampling periods. Points indicate the mean and standard error of the hybrid index at each sampling locality, distance starts at 0 km at sampling locality 1, and shading represents the 95% bootstrap confidence interval. (**B**) Cline centers with 95% confidence intervals for the geographic clines estimated from the hybrid index and separately for the six plumage traits. Full model details are available in Table S6.

**FIGURE 3.**
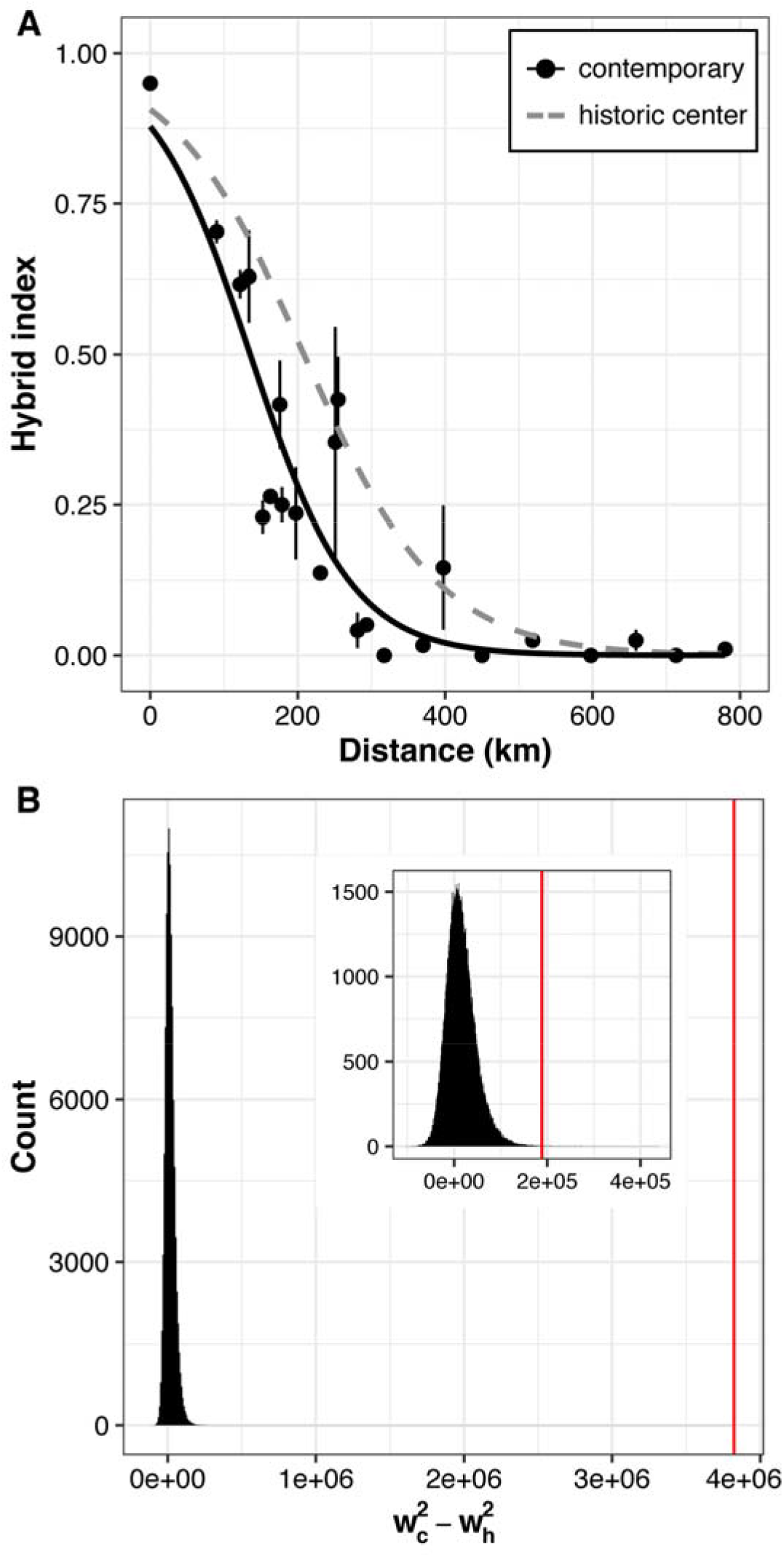
(**A**) Comparison of the true contemporary geographic cline for hybrid index (black line, circles) to a contemporary cline with the center fixed to the estimate of the historic center (gray dashed line). ΔAIC indicates the true cline is significantly better, further supporting the difference in cline centers between the historic and contemporary sampling periods. (**B**) The bootstrap distribution of 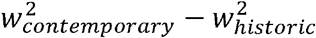 of the hybrid index cline is significantly different from the expectation (red line) under the neutral diffusion model for both realistic (main plot) and conservative (inset plot) values of dispersal and generation time, suggesting the hybrid zone has not widened to the extent expected under neutral diffusion.

The hybrid zone movement we identify is robust to various checks of the dataset (Table S7): cline models made separately for males and females are similar to the full model in both sampling periods (Fig. S3); contemporary localities sampled on the North Platte do not overly influence the full cline model (Fig. S4); and the fact that sampling localities are not completely identical across both sampling periods does not overly influence the full model in either period (Fig. S5). Looking only at localities that are directly resampled in both periods, we find the expected pattern of shifts in the hybrid index from higher scores (more red-shafted) in the historic samples to lower scores (more yellow-shafted) in the contemporary samples in the central part of the hybrid zone as would happen with a westward shift of the hybrid zone (Fig. S6, Table S8).

## Discussion

The hybrid zone between red-shafted and yellow-shafted flickers in the Great Plains of North America was an important study system in the early development of ideas about hybrid zone dynamics (e.g., Moore and Buchanan 1985; Moore and Price 1993). In this study, we compared historic (1955-1957) and contemporary (2015-2018) samplings of an identical transect across the flicker hybrid zone to assess changes over the past ∼60 years. We focused on patterns in six plumage characteristics using a categorical plumage scoring approach that we validated with independent multispectral photography (Table S4, S5; Fig. S1).

We detected a significant westward shift of the hybrid zone center of ∼73 km towards the range of the red-shafted flicker (Fig. 2A, 3A). In the historic sampling period, the cline center was ∼208 km east of the start of the transect (between localities 11 and 12 in Fig. 1), while in the contemporary sampling period, the center shifted to ∼135 km east of the start of the transect (near localities 5 and 6). Our estimation of the center of the cline in the historic sampling period matches previous work using the same samples with different methods (Short 1965) and different samples and methods (Moore and Buchanan 1985). Moreover, we conducted a variety of analytic checks of the dataset in both the historic and contemporary sampling periods, and these provided further support for the contemporary movement of the hybrid zone that we identified (Fig. S3-6). This movement differs greatly from previous work done in the same region that instead found geographic stability when comparing samples between the periods of 1889-1968 and 1981-1982 (Moore and Buchanan 1985). Thus, it seems likely that the flicker hybrid zone moved in the latter ∼35 years between our historic and contemporary sampling points (i.e., after the study by Moore and Buchanan 1985), which suggests the rate of westward movement has been quite rapid: ∼2.1 km/year since the early 1980s.

Despite this rapid movement, the flicker hybrid zone has remained remarkably narrow between our two sampling periods. We did not find evidence of significant changes in the width of the flicker hybrid zone over our ∼60-year sampling period (Table S6) and our analytical checks of the dataset further underscore this stability in width (Table S7). Using an approach that takes advantage of repeat sampling of the same transect (Wang et al. 2019), we were additionally able to directly refute the expectations of neutral diffusion (Fig. 3B; Barton and Hewitt 1985). These results support some selective force preventing changes in the width of the hybrid zone, as was similarly found in Moore and Buchanan (1985). However, the lack of evidence for fitness consequences of hybridization despite intensive field studies within the hybrid zone (e.g., Moore and Koenig 1986; Wiebe and Bortolotti 2002; Flockhart and Wiebe 2008, 2009) means the hybrid zone is not well-explained as a tension zone model (Barton and Hewitt 1985, 1989).

Instead, the flicker hybrid zone may be better described by an environmental selection gradient model where hybrids have higher fitness than parentals within the hybrid zone (Moore 1977; Moore and Price 1993). Under this model, if there is ecological change that moves the geographic area where hybrids have higher fitness than parentals, there can be hybrid zone movement to track this change. This movement could occur without associated changes in width as long as the geographic area of higher hybrid fitness moves without expanding or narrowing. Thus, the rapid movement of the flicker hybrid zone since the early 1980s may be tied to changes in the environment. Intriguingly, similar westward movements have been documented in two of the other four major Great Plains avian hybrid zones in the same geographic region. The hybrid zone between lazuli (*Passerina amoena*) and indigo (*P. cyanea*) buntings has shifted westward ∼100 km (Carling and Zuckerberg 2011); and the hybrid zone between Baltimore (*Icterus galbula*) and Bullock’s (*I. bullockii*) orioles has shifted westward 41-73 km (Corbin and Sibley 1977; Walsh et al. 2020). Westward movements repeated across three separate Great Plains hybrid zones suggest these movements are not the result of sampling error. Moreover, the similarity in magnitude between the movement of the flicker, bunting, and oriole hybrid zones over similar sampling periods is suggestive of a shared driver. Niche modelling has suggested that differences in temperature influence the locations of the avian Great Plains hybrid zones (Swenson 2006). In both buntings and orioles, the western species of the hybridizing pair is adapted to hotter, drier environments, while the eastern species is adapted to wetter environments (Rising 1969; Carling and Thomassen 2012). Although, to date, similar adaptive differences have not been found between the two flickers, the movement of these hybrid zones could be the result of environmental changes in the Great Plains that has shifted the location of favorable habitat for the eastern species.

For instance, land management and irrigation practices over the last century have influenced the flow of the Platte River and subsequently, the extent of surrounding woodland riparian corridors such that areas that were historically treeless or sparsely wooded now support mature cottonwoods and willows (Johnson 1994, 1997; Horn et al. 2012). Flickers rely on the large trees in these riparian corridors for nesting cavities, so this woody encroachment may have allowed for more dispersal of breeding yellow-shafted flickers into the Great Plains and a resulting westward shift of the hybrid zone center. It’s also possible, however, that climate change may be causing directional movement of the Great Plains ecotone itself, as has been demonstrated in other ecotones (Smith and Goetz 2021). Because the ecotone is oriented on the east-west axis, any movement of the ecotone would also occur along this axis. Thus, if the flicker hybrid zone tracks aspects of the ecotone, then movement would occur in the eastern or western direction as we have observed. Unfortunately, until we can characterize the selective forces maintaining the hybrid zone, it will remain difficult to assess whether changes in land management, climate, or a combination of factors are most influential.

We expect the patterns we identified for the hybrid zone center and width of plumage characteristics to be representative of patterns at the genotypic level for flickers. Previous work has demonstrated the extremely low levels of genomic divergence that exist between red-shafted and yellow-shafted flickers away from the hybrid zone (Aguillon et al. 2018, 2021): 780 fixed differences across ∼7.25 million markers. As the few existing differentiated regions of the genome underly their coloration differences (Aguillon et al. 2021), we believe our results using plumage scoring additionally represent a good proxy for the patterns at the genomic level.

Our results underscore the importance of biological collections. We identified a significant westward movement in the long-studied flicker hybrid zone that may have gone unnoticed without repeat sampling efforts—something that is difficult to accomplish without the long memory of collections. We hope that in another 60 years additional sampling of this transect along the flicker hybrid zone will be undertaken and provide further discoveries about the evolutionary process and the maintenance of hybrid zones over time. In the meantime, future work should investigate the cause of this westward movement and its relationship to environmental changes.

## AUTHOR CONTRIBUTIONS

S.M.A. conceived the study, analyzed the data, and wrote the original draft of the manuscript. S.M.A. and V.G.R. collected the data and revised the final manuscript.

## ACKNOWLEDGMENTS

We greatly appreciate the insights, guidance, and resources provided by I. J. Lovette for this project. We thank R. A. Ligon for generously providing the photography equipment (and the know-how) necessary to take the multispectral images and for providing helpful feedback on data processing. M. Bishop provided illustrations of the flickers. We thank B. Mims, N. A. Kramer, and T. Brooks for providing assistance in the field and G.-F. Siegmund for input on data analysis. The Schumer Lab, S. Singhal, J. Walsh, J. B. Searle, L. Campagna, and M. N. Vitousek provided helpful comments on earlier versions of this manuscript. This work was supported by the Cornell Lab of Ornithology Athena Fund, the Garden Club of America Frances M. Peacock Scholarship, and the Cornell University EEB Betty Miller Francis and Paul P. Feeny Graduate Student Research Funds (all to S.M.A.). S.M.A. was supported by the US National Science Foundation Graduate Research Fellowship Program (DGE-1144153) and the AAUW American Dissertation Fellowship. Science is only moved forward by standing on the shoulders of those who came before, so we would like to acknowledge the ground-breaking efforts undertaken by Lester L. Short to collect and study flickers in the 1950s.

## DATA ARCHIVING

Datafiles will be made available on the Dryad Digital Repository upon publication and all scripts will be available on GitHub.

## Footnotes

1 The subspecific epithet of the red-shafted flicker is based on a term that is an extreme racial slur against Black Africans, particularly in South Africa. We include the official scientific name here, but purposefully refer to the flickers elsewhere only by their common names. Changing the name to *Colaptes auratus lathami* has been proposed (Aguillon and Lovette 2019), but this has not been officially accepted.

## Notes

https://github.com/stepfanie-aguillon/flicker-HZ-movement

### Competing Interest Statement

The authors have declared no competing interest.

