## Supplement for "Revisiting a classic hybrid zone: rapid movement of the northern flicker hybrid zone in contemporary times"

### Electronic Supplementary Material

**Text S1. Detailed methods for multispectral photography**

We collected multispectral images of all contemporary CUMV flicker specimens (Table S2), to obtain a more quantitative assessment of phenotypic traits following (Ligon et al. 2018). We took RAW format images under standardized conditions using a Canon 7D camera (Tokyo, Japan) with full-spectrum quartz conversion and fitted with a Novoflex Noflexar 35 mm lens. Specimens were illuminated by two eyeColor arc lamps (Iwasaki, Tokyo, Japan) that simulate CIE-recommended daylight (D65) and were diffused through 0.5 mm polytetrafluoroethylene sheets. The lamps include a UV-blocking coating, which was removed prior to image collection. To capture the important aspects of flicker plumage, each specimen was photographed from three viewing angles: ventral (flat on its back), dorsal (flat on its belly), and lateral (on its left side). Additionally, for each viewing angle, we used filters (Baader, Mammendorf, Germany) to take two photographs: one capturing visible light between 400-700 nm and one capturing only UV light between 300-400 nm. All specimens were photographed against a blank white background and included size and color standards. For visible light photographs, we used a shutter speed of 1/6” and ISO of 400, and for UV photographs those parameters were increased to 4” and 3200, respectively.

Visible and UV photographs were used to create standardized multispectral image files for each specimen and viewing angle using the Multispectral Image Calibration and Analysis Toolbox (micaToolbox; Troscianko and Stevens 2015) in ImageJ (Schneider et al. 2012). Standardized multispectral images combine, equalize, and linearize the different color channels from the two photographs (Stevens et al. 2007). We identified regions of interest (ROI) within each flicker as follows: throat and shaft with the ventral viewing angle; crown and nuchal patch with the dorsal viewing angle; and ear coverts and malar stripe with the lateral viewing angle. We then used the micaToolbox to estimate the color sensitivity of our camera/lens combination and generate a custom mapping function to convert colors in the image to stimulation values corresponding to an avian visual space (the Eurasian blue tit, *Cyanistes caeruleus*). We used the Batch Multispectral Image Analysis option in the micaToolbox to output values for each color channel (long wave, medium wave, short wave, UV) and luminance within each ROI, as well as the overall area of the ROI (important only for the nuchal patch).

**TABLE S1.** Details on the scoring of six plumage trait differences between red-shafted and yellow-shafted flickers. This method is slightly adapted from Short (1965).

| Plumage score | Description |
|---|---|
| *Crown color* | |
| 0 | Gray, as in yellow-shafted |
| 1 | Gray with brown traces in forehead and crown |
| 2 | Mixed gray and brown (crown half brown with more gray on hind neck) |
| 3 | Crown brown with hind neck gray toward back |
| 4 | Brown, as in red-shafted |
| *Ear covert color* | |
| 0 | Tan, as in yellow-shafted |
| 1 | Tan with gray traces |
| 2 | Mixed gray and tan |
| 3 | Gray with tan traces (especially below eye) |
| 4 | Gray, as in red-shafted |
| *Malar stripe color (males only)* | |
| 0 | Black, as in yellow-shafted |
| 1 | Black with <20% red |
| 2 | Mixed black and red |
| 3 | Red with <20% black |
| 4 | Red, as in red-shafted |
| *Nuchal patch presence* | |
| 0 | Present and broad, as in yellow-shafted |
| 1 | Present and restricted in width (less than one-half of normal width) |
| 2 | Present and broken in one or more places |
| 3 | Traces present, usually at sides of nape |
| 4 | Absent, as in red-shafted |
| *Shaft color* | |
| 0 | Bright yellow, as in yellow-shafted |
| 1 | Yellow-orange |
| 2 | Orange |
| 3 | Red-orange |
| 4 | Deep salmon red, as in red-shafted |
| *Throat color* | |
| 0 | Tan, as in yellow-shafted |
| 1 | Tan with gray traces (usually on lower throat) |
| 2 | Mixed gray and tan |
| 3 | Gray with tan traces (usually near chin) |
| 4 | Gray, as in red-shafted |

**TABLE S2.** Information on samples included in this study. Locality ID/Name correspond to values in Fig. 1 and Table S3. Plumage scores range from 0 (yellow-shafted) to 4 (red-shafted). The overall hybrid index has been transformed to range from 0 to 1 to allow comparisons between the sexes. *Table included as separate excel file*

**TABLE S3.** Sampling localities used in the geographic cline analyses and shown in the inset map in Fig. 1.

| Locality ID | Locality Name | Latitude | Longitude | Distance | Sampling Period |
|---|---|---|---|---|---|
| 1 | Western | 40.8830559 | -105.51687 | 0 | contemporary |
| 2 | Greeley | 40.346603 | -104.82279 | 58.4 | historic |
| 3 | Kersey | 40.3693671 | -104.44526 | 90.1 | contemporary |
| 4 | Orchard | 40.351842 | -104.06838 | 121.8 | historic and contemporary |
| 5 | W Fort Morgan | 40.267982 | -103.94052 | 132.6 | historic |
| 6 | Morrill | 41.9594945 | -103.92463 | 133.9 | contemporary |
| 7 | E Fort Morgan | 40.2825194 | -103.70253 | 152.6 | contemporary |
| 8 | Brush | 40.3272657 | -103.57831 | 163.0 | contemporary |
| 9 | Minatare | 41.8984992 | -103.42772 | 175.7 | contemporary |
| 10 | Merino | 40.4047694 | -103.39182 | 178.7 | contemporary |
| 11 | Bridgeport | 41.6895056 | -103.16821 | 197.5 | contemporary |
| 12 | Crook | 40.8444414 | -102.77278 | 230.7 | historic and contemporary |
| 13 | Lisco | 41.4499137 | -102.53066 | 251.1 | contemporary |
| 14 | Sedgwick | 40.932974 | -102.48678 | 254.8 | contemporary |
| 15 | Big Springs | 40.9935125 | -102.17377 | 281.1 | historic and contemporary |
| 16 | Lewellen | 41.2998817 | -102.02722 | 293.4 | contemporary |
| 17 | Ogallala | 41.22806 | -101.74345 | 317.3 | contemporary |
| 18 | Sutherland | 41.1451911 | -101.11715 | 369.9 | historic and contemporary |
| 19 | North Platte | 41.17314 | -100.78951 | 397.5 | contemporary |
| 20 | Halsey | 41.903358 | -100.31039 | 437.7 | historic |
| 21 | Gothenberg | 40.9061571 | -100.16424 | 450.0 | historic and contemporary |
| 22 | Elm Creek | 40.686218 | -99.347335 | 518.7 | historic and contemporary |
| 23 | Burwell | 41.756348 | -99.282391 | 524.1 | historic |
| 24 | Grand Island | 40.853711 | -98.406833 | 597.7 | historic and contemporary |
| 25 | Silver Creek | 41.2795413 | -97.680686 | 658.7 | historic and contemporary |
| 26 | Schuyler | 41.4080137 | -97.031092 | 713.3 | historic and contemporary |
| 27 | Eastern | 41.3335076 | -96.238837 | 779.8 | historic and contemporary |

**TABLE S4.** Overall results from multiple regressions comparing the plumage score with values from the multispectral photography for each plumage trait. For the nuchal patch, we compared the plumage score to the patch area in the image. For all other traits, we compared the plumage score to the values for the four color channels and luminance.

| Trait | Adjusted $R^2$ | p-value |
|---|---|---|
| crown | 0.398 | $3.109 \times 10^{-9}$ |
| ear coverts | 0.6812 | $< 2.2 \times 10^{-16}$ |
| malar stripe | 0.8718 | $< 2.2 \times 10^{-16}$ |
| nuchal patch | 0.6538 | $< 2.2 \times 10^{-16}$ |
| shaft | 0.9099 | $< 2.2 \times 10^{-16}$ |
| throat | 0.8541 | $< 2.2 \times 10^{-16}$ |

**TABLE S5.** Detailed model results from multiple regressions comparing the plumage score with values from the multispectral photography. Overall model results are in Table S4 and visualized in Fig. S6.

| Trait | Value | Estimate | SE | t | p-value |
|---|---|---|---|---|---|
| crown | intercept | 20.88 | 10.74 | 1.944 | 0.0553 |
| crown | lwMean | -1448.00 | 1515.57 | -0.955 | 0.3421 |
| crown | mwMean | -3935.41 | 3219.34 | -1.222 | 0.2250 |
| crown | swMean | -1021.14 | 978.95 | -1.043 | 0.2999 |
| crown | uvMean | 34.08 | 91.56 | 0.372 | 0.7107 |
| crown | lumMean | 6273.81 | 5591.50 | 1.122 | 0.2650 |
| ear coverts | intercept | 10.66 | 7.52 | 1.417 | 0.1600 |
| ear coverts | lwMean | -1405.15 | 1162.72 | -1.209 | 0.2300 |
| ear coverts | mwMean | -2749.38 | 2376.04 | -1.157 | 0.2510 |
| ear coverts | swMean | -776.49 | 757.36 | -1.025 | 0.3080 |
| ear coverts | uvMean | 40.74 | 65.10 | 0.626 | 0.5330 |
| ear coverts | lumMean | 4882.61 | 4213.41 | 1.159 | 0.2500 |
| malar stripe | intercept | 10.63 | 9.19 | 1.156 | 0.2530 |
| malar stripe | lwMean | -1418.97 | 1434.26 | -0.989 | 0.3270 |
| malar stripe | mwMean | -2958.76 | 2890.22 | -1.024 | 0.3110 |
| malar stripe | swMean | -929.03 | 933.33 | -0.995 | 0.3240 |
| malar stripe | uvMean | 44.26 | 75.16 | 0.589 | 0.5580 |
| malar stripe | lumMean | 5228.49 | 5160.38 | 1.013 | 0.3160 |
| nuchal patch | intercept | 2.91 | 0.18 | 16.08 | $< 2 \times 10^{-16}$ |
| nuchal patch | area | 0.00 | 0.00 | -13.00 | $< 2 \times 10^{-16}$ |
| shaft | intercept | -3.20 | 1.76 | -1.820 | 0.0723 |
| shaft | lwMean | 798.97 | 251.95 | 3.171 | 0.0021 |
| shaft | mwMean | 1450.13 | 508.49 | 2.852 | 0.0055 |
| shaft | swMean | 535.96 | 159.38 | 3.363 | 0.0012 |
| shaft | uvMean | -34.76 | 7.80 | -4.457 | 0.0000 |
| shaft | lumMean | -2725.39 | 908.22 | -3.001 | 0.0035 |
| throat | intercept | 7.41 | 3.54 | 2.097 | 0.0390 |
| throat | lwMean | -930.08 | 535.09 | -1.738 | 0.0858 |
| throat | mwMean | -1550.25 | 1096.90 | -1.413 | 0.1613 |
| throat | swMean | -588.43 | 353.36 | -1.665 | 0.0996 |
| throat | uvMean | 71.33 | 35.49 | 2.010 | 0.0477 |
| throat | lumMean | 2994.04 | 1941.64 | 1.542 | 0.1268 |

**TABLE S6.** Model results for the geographic clines from plumage scoring in the historic and contemporary sampling periods. Clines are visualized in Fig. 2 and S2.

| Trait | Center | Center 95% CI | Width | Width 95% CI |
|---|---|---|---|---|
| *Historic sampling period* | | | | |
| hybrid index | 208.3763 | 190.0316, 226.8876 | 251.1938 | 188.8206, 328.1764 |
| crown | 277.0663 | 206.6787, 351.9533 | 462.8924 | 149.3879, 826.9237 |
| ear coverts | 153.8192 | 144.0973, 163.6192 | 267.7168 | 232.0703, 307.8535 |
| malar stripe | 218.0839 | 203.8221, 231.8209 | 175.5512 | 122.5301, 227.6591 |
| nuchal patch | 228.9754 | 205.2576, 251.3624 | 173.7698 | 62.07621, 268.8413 |
| shaft | 207.8027 | 184.1068, 231.5219* | 211.4893 | 134.074, 294.1099* |
| throat | 191.2107 | 173.0066, 209.2131 | 337.3793 | 275.8701, 409.57 |
| *Contemporary sampling period* | | | | |
| hybrid index | 135.6074 | 108.7064, 156.7343 | 274.5806 | 179.1994, 408.213 |
| crown | 114.2603 | 74.6308, 140.5735 | 237.7157 | 109.6608, 422.9463 |
| ear coverts | 128.313 | 99.8488, 148.052 | 179.0388 | 103.3117, 303.813 |
| malar stripe | 186.9072 | 132.0964, 235.1586 | 332.3642 | 173.667, 677.4881 |
| nuchal patch | 115.918 | 77.6389, 141.2332 | 263.3228 | 147.1721, 429.0008 |
| shaft | 116.5658 | 38.8284, 158.1011 | 300.6879 | 28.15649, 670.1835 |
| throat | 179.5272 | 151.5103, 205.1583 | 261.3114 | 162.6929, 427.2188 |

* Confidence intervals based on bootstrapping.

**TABLE S7.** Analytical checks of the geographic cline model results for the overall hybrid index. The dataset has been separated by sex (Fig. S3), filtered to remove the sampling localities from the North Platte (Fig. S4), and subset to include only localities with repeat sampling (Fig. S5) to validate the full model results presented in Fig. 2 and Table S6.

| Trait | Center | Center 95% CI | Width | Width 95% CI |
|---|---|---|---|---|
| *Historic sampling period* | | | | |
| **full model** | **208.3763** | **190.0316, 226.8876** | **251.1938** | **188.8206, 328.1764** |
| females only | 214.5113 | 195.1848, 233.3106 | 202.6322 | 133.7830, 279.6673 |
| males only | 205.3811 | 186.5932, 224.6111 | 286.8929 | 219.7600, 368.7244 |
| repeat sampling | 203.8806 | 175.6011, 228.0885 | 253.7850 | 163.9867, 377.1234 |
| *Contemporary sampling period* | | | | |
| **full model** | **135.6074** | **108.7064, 156.7343** | **274.5806** | **179.1994, 408.213** |
| females only | 117.2019 | 12.3450, 159.4695 | 351.2353 | 164.2160, 825.3515 |
| males only | 138.4291 | 101.1033, 165.9318 | 278.0852 | 151.4056, 472.1344 |
| repeat sampling | 143.9636 | 138.4320, 149.4619 | 185.9734 | 165.7705, 207.5685 |
| South Platte | 126.6724 | 96.3382, 151.6241 | 246.5787 | 121.5106, 423.3045 |

**TABLE S8.** Two-sided Wilcoxon rank-sum tests comparing the hybrid index between the historic and contemporary sampling periods at localities that were directly resampled. Results are visualized in Fig. S6.

| Locality ID | Locality Name | N | W | p-value |
|---|---|---|---|---|
| 4 | Orchard | 9 | 2 | 0.184 |
| 12 | Crook | 24 | 27 | 0.0164 |
| 15 | Big Springs | 9 | 2.5 | 0.2356 |
| 18 | Sutherland | 32 | 53.5 | 0.4295 |
| 21 | Gothenberg | 38 | 32.5 | 0.0233 |
| 22 | Elm Creek | 28 | 51.5 | 0.7242 |
| 24 | Grand Island | 43 | 8 | 0.2952 |
| 25 | Silver Creek | 31 | 28 | 0.9648 |
| 26 | Schuyler | 40 | 46 | 0.1656 |
| 27 | Eastern | 5 | 2.5 | 1 |

**FIGURE S1.** Relationships between plumage score and image parameters from the multispectral photography. Plumage scores are summarized with box plots underneath the data (colored points). For the nuchal patch, we compared the plumage score to the patch area in the image. For all other traits, we compared the plumage score to the values for the four color channels and luminance. The analysis was run as a multiple regression (Table S4 and S5), but we show here the sum of all of the photo values.

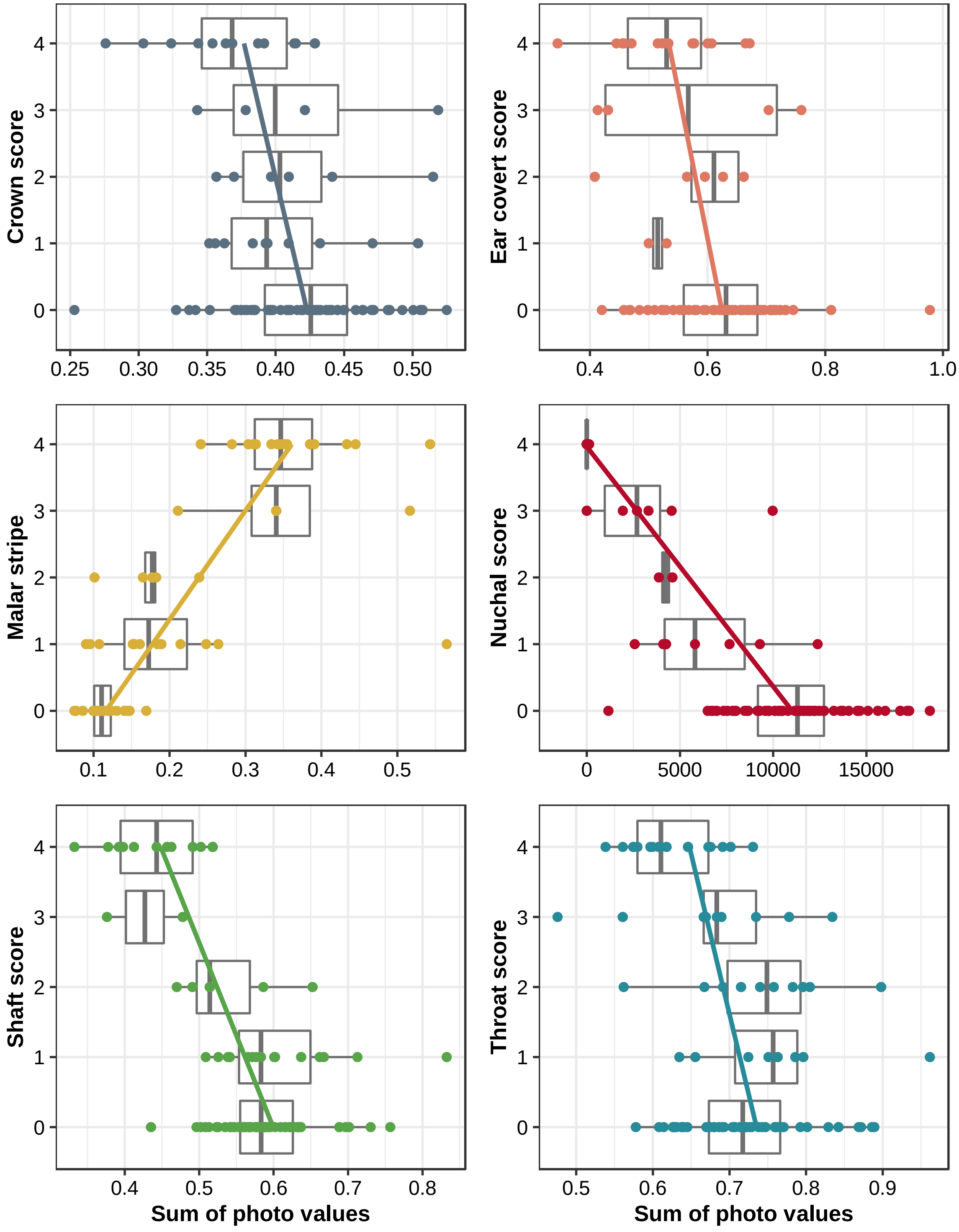

**FIGURE S2.** Geographic clines of the six plumage traits for the (**A**) historic and (**B**) contemporary sampling periods. Traits are indicated by different colors and points indicate the average trait score at each sampling locality (jittered for visualization). Corresponding model outputs are available in Table S6.

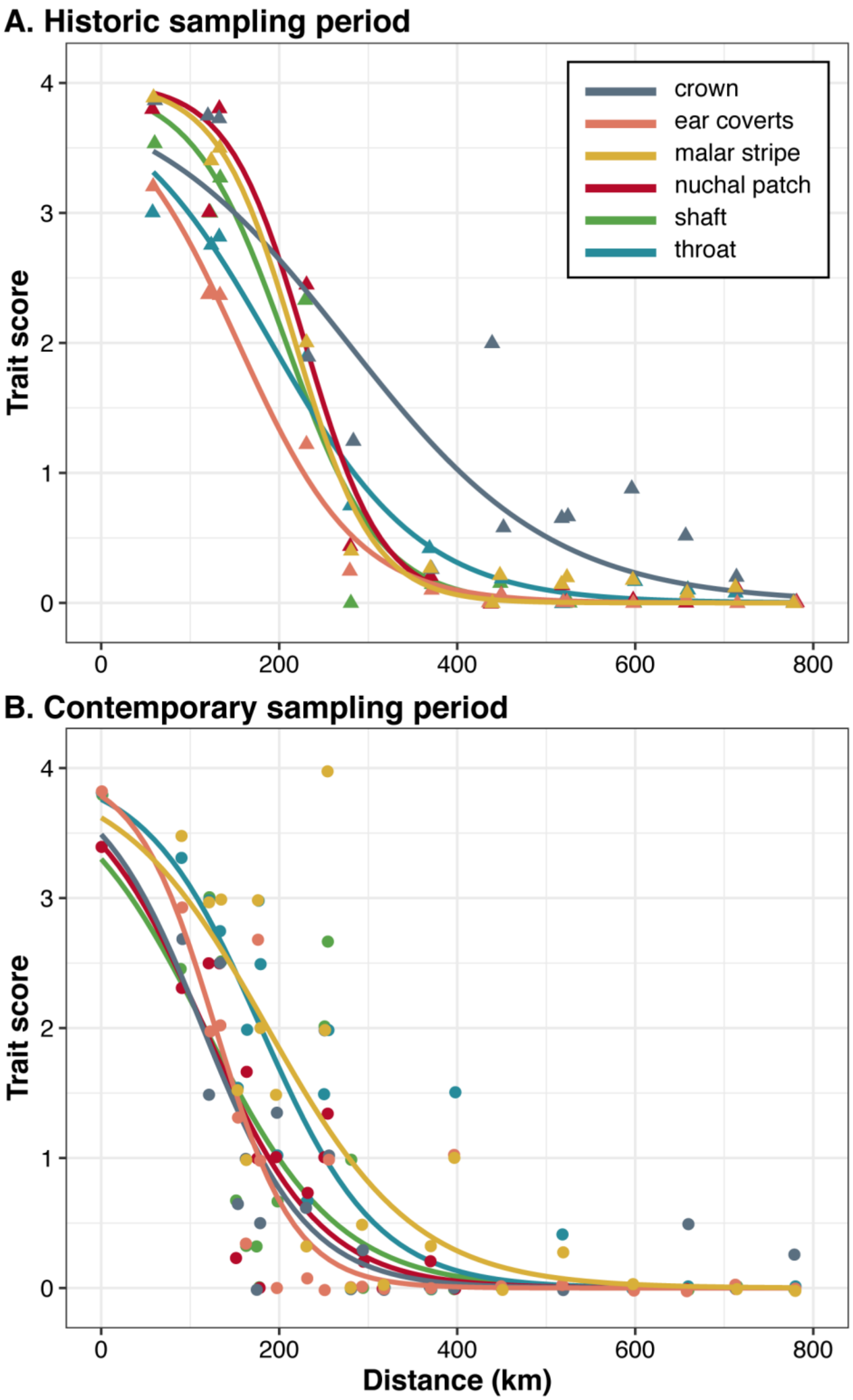

**FIGURE S3.** Geographic clines of the hybrid index for the (**A**) historic and (**B**) contemporary sampling periods with the dataset separated by sex (females = reds, males = blues). Full model results are shown with solid lines and shading represents the 95% bootstrap confidence interval. Corresponding model outputs are available in Table S7.

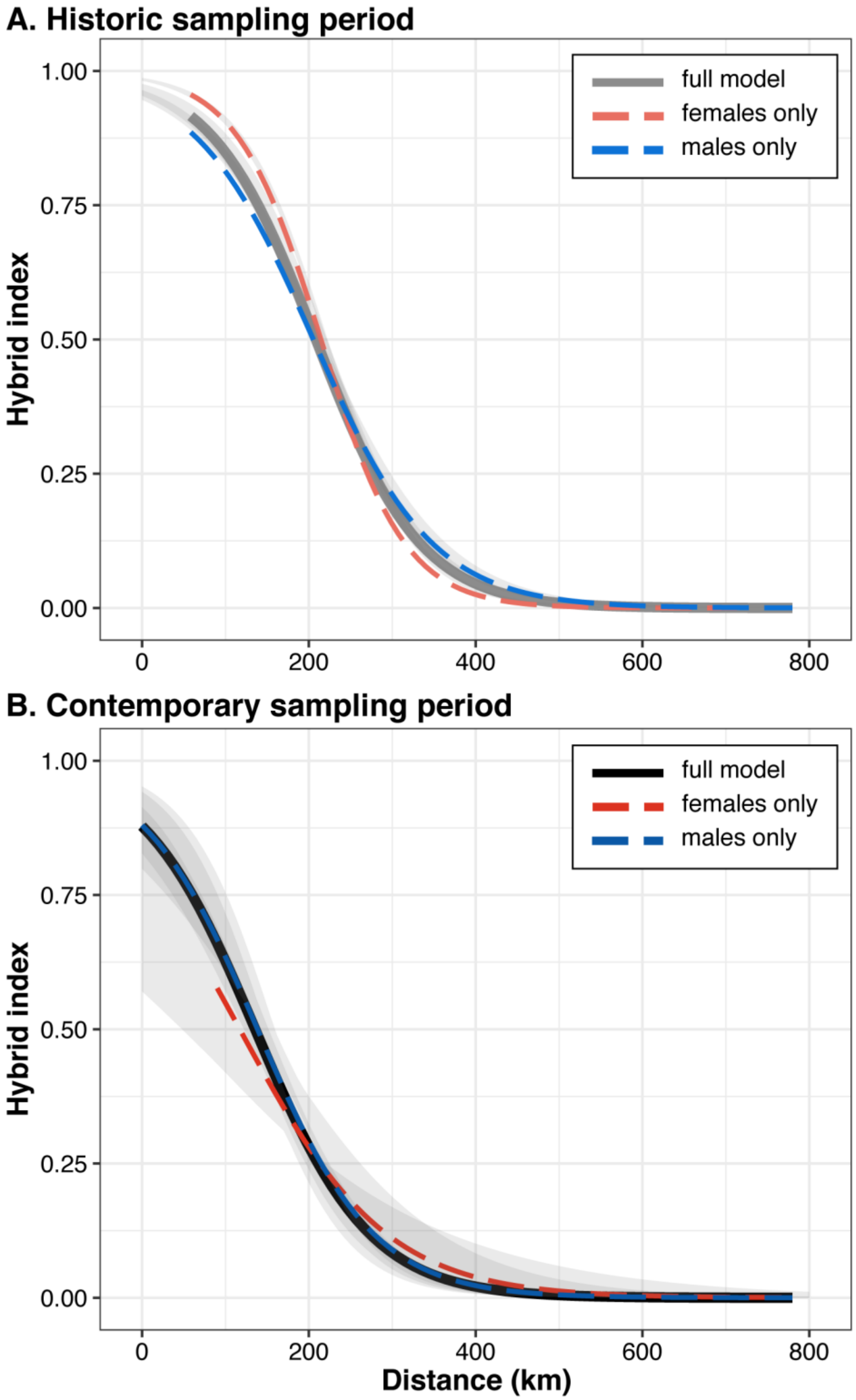

**FIGURE S4.** Geographic cline of the hybrid index for the contemporary sampling period with the data filtered to remove the sampling localities from the North Platte (red, dashed line). Full model results are shown with the solid black line, the removed localities are shown with open circles, and shading represents the 95% bootstrap confidence interval. Corresponding model outputs are available in Table S7.

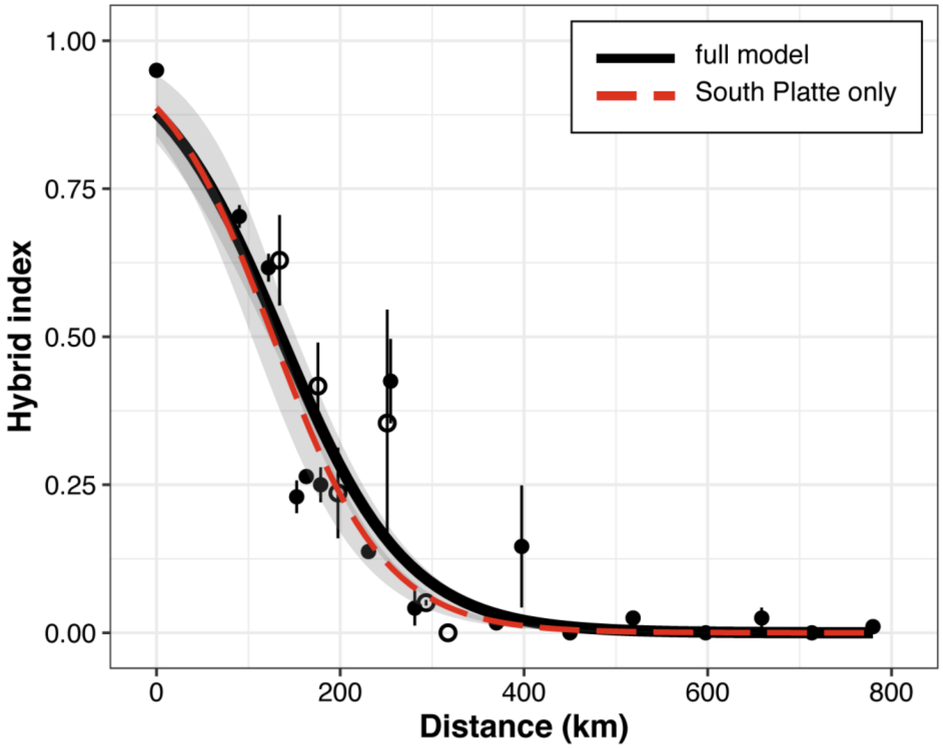

**FIGURE S5.** Geographic clines of the hybrid index for the historic (pink, dashed line) and contemporary (red, dashed line) sampling periods with the dataset subset to include only the 10 localities sampled during both time periods. Full model results are shown with solid lines and shading represents the 95% bootstrap confidence interval. Corresponding model outputs are available in Table S7.

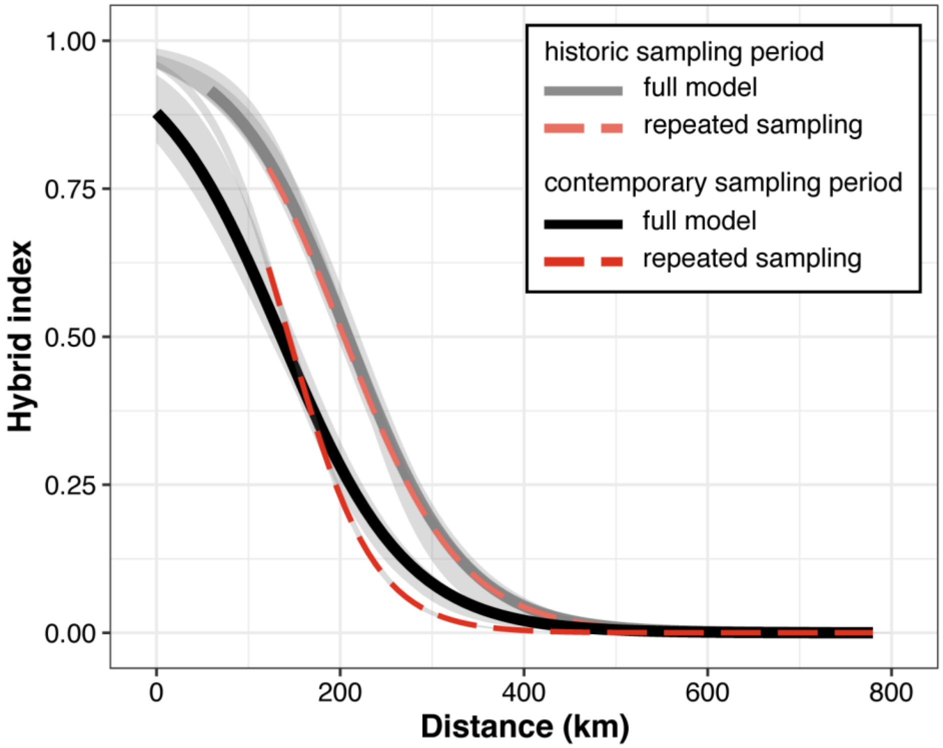

**FIGURE S6.** Distribution of the hybrid index at the 10 localities sampled during both the historic (gray) and contemporary (black) sampling periods. The movement of the hybrid zone is apparent in the shift in the hybrid index towards lower scores in the contemporary samples (more yellow-shafted) at the localities in the more central part of the hybrid zone (upper row). Due to small sample sizes in the contemporary samples, this trend is only significant at locality 12 and 21. Full statistical results are shown in Table S8.

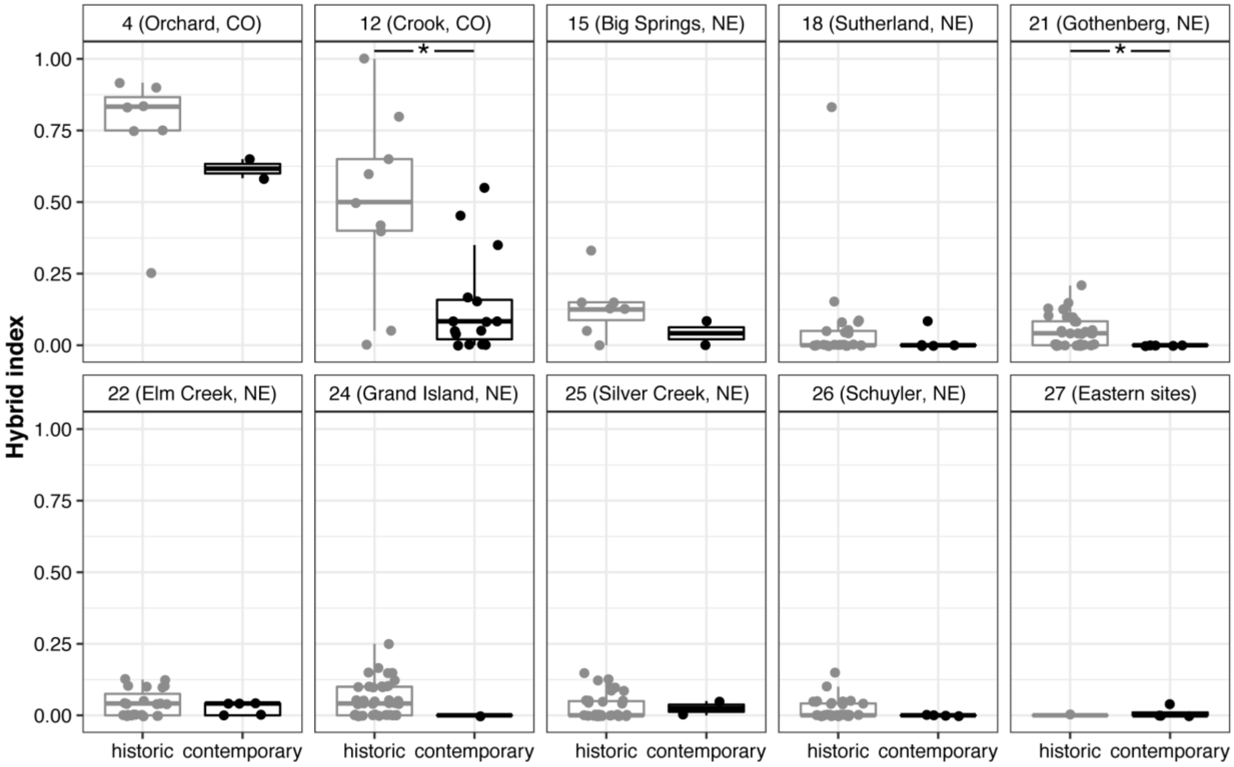
